# CVRMS: Cross-validated Rank-based Marker Selection for Genome-wide Prediction of Low Heritability

**DOI:** 10.1101/756130

**Authors:** Seongmun Jeong, Jae-Yoon Kim, Namshin Kim

## Abstract

CVRMS is an R package designed to extract marker subsets from repeated rank-based marker datasets generated from genome-wide association studies or marker effects for genome-wide prediction (https://github.com/lovemun/CVRMS). CVRMS provides an optimized genome-wide biomarker set with the best predictability of phenotype by implemented ridge regression using genetic information. Applying our method to human, animal, and plant datasets with wide heritability (zero to one), we selected hundreds to thousands of biomarkers for precise prediction.

---

Genomic prediction (GP) (Meuwissen et al., 2001) is a potent tool in plant and animal breeding and in the prediction of complex traits and disease risks in humans. Several GP methods, such as the method of marker effect (Meuwissen., 2001; Habier et al., 2011; Gianola, 2013) and genomic best linear unbiased prediction (GBLUP) (VanRaden, 2008), have been proposed over the years.

A traditional statistical prediction approach is that large numbers of predictor variables, such as SNPs, are regressed on phenotypes. Recently, as SNP density and sample size have increased and analysis technology has improved, a precise and efficient method for making such predictions has become necessary. GP is a method to predict the trait of interest based on the genomic-based estimated value (GBEV) predicted from the genome-scale dataset (Hayes et al., 2009). GBEV is calculated based on the effect of markers from the training population using statistical modeling or machine learning methods. Various models, including an improved statistical method and those based on machine learning and deep learning, have been developed for GP.

Ridge-regression best linear unbiased prediction (RR-BLUP) (Meuwissen et al., 2001) was the first to be adopted and assumes that all effects of the markers are normally distributed and have equal variance. Although improved methods such as the genomic BLUP (Habier et al., 2007) and the linear genomic BLUP kernel (Gianola et al., 2006) have been subsequently presented, RR-BLUP is still used frequently due to additional computational issues, including the genomic relationship matrix or the pairwise distance matrix between samples.

Many essential plants, animal, and human traits and diseases are moderately heritable, implying that differences in DNA sequences can attribute a large portion of the differences between individuals in phenotypes. For those heritable traits, prediction of the phenotype or risk using DNA can usually reach moderately high prediction accuracy (de los Campos et al., 2018). Thus, GP tends to increase prediction accuracy with higher heritability (Zhang et al., 2017). In general, high-heritability traits, including flowering time in plants and human height, have already been studied (de Oliveira et al., 2018 and Lello et al., 2018). Among the same traits, heritability is low according to the characteristics of the population (for example, days to silk in maize), and there are traits with low heritability regardless of population (for example, human complex traits). In this case, genomic prediction of these traits is very difficult, and there are only a few cases where the accuracy of the model is improved (Zhang et al., 2019, Guarini et al., 2019, Ward et al., 2019, crop science).

Human genomic prediction studies (Khera et al., 2019, Mavaddat et al., 2019, and Lamri et al., 2019) have been mainly based on the polygenic risk score (PRS), but this method relies on known markers associated with the trait, which tend to have low allele frequency. In plant breeding programs, marker-assisted selection (MAS), which is an indirect selection based on a marker linked to a trait of interest, is a commonly used method for genomic prediction. Another common problem of genomic prediction is low heritability. Complex traits often have different heritabilities, for example, longevity (0.25, Herskind et al., 1996), body mass index (0.05~0.9, Allison et al., 1996) and height (~0.8, Yang et al., 2015).

Since the selected SNPs may vary depending on the phenotypes, it is necessary to set specific SNPs for each target characteristic (Sousa et al., 2019). This process is often called feature selection, in which subsets of available features are selected for application in prediction models. The best subset of features contains the least number of dimensions that most contribute to prediction accuracy (Guyon et al., 2003). Several studies (Bermingham et al., 2015, Li et al., 2018, and Lenz et al., 2017) have performed marker selection by selecting most significant SNPs through the result of a genome-wide association study (GWAS) or marker effect analysis, but selecting a marker in this way only considers the relationship between the marker and the trait, not the correlation among the markers.

In this study, we developed an implemented marker selection method, called CVRMS, to improve the accuracy of genomic prediction by selecting a minimized marker subset considering the low heritability problem and the correlation between markers. Its algorithm is simple but powerful, so it can perform marker selection even when the heritability of a given trait is relatively low.

We applied CVRMS to select a marker subset for human height and a flowering time of two plant datasets. We used 413 diverse rice (*O. Sativa*) varieties from 82 countries using a high-quality, custom-designed, 44,100 oligonucleotide genotyping array and the flowering times from two regions (Arkansas, U.S. and Faridpur, Bangladesh) (Zhao et al., 2011). We calculated p-values to conduct genomewide association analysis using GAPIT (Tang et al., 2016) for the rice dataset. Distribution plot (manhattan plot) of p-values with log10 scale was shown in supplementary figure 1. CVRMS with 5-fold cross-validation, ten minimum markers, 300 maximum markers, 0.001 delta, and 0.9 accuracy cutoff was used to select marker subsets from the genotype, phenotype datasets and the GAPIT GWAS results. 67 and 88 markers were selected and have the prediction accuracy (pearson correlation coefficient) with 0.9028 (0.6332 for whole markers and 0.8695 for validation set) and 0.9018 (0.4425 for whole markers and 0.8329 for validation set) for the flowering time in Arkansas and Faridpur, respectively (Supplementary Table 1).

In the case of the human height dataset, we obtained data from the Korean population-based cohort dataset (8,840 individuals) of the Korean Genome Epidemiology Study (KoGES) with Affymmetrix SNP Array 5.0. Imputation was conducted by impute2 with a 1000 genome reference panel, and 1,818,708 SNPs were used for genome-wide association studies with PLINK (Chang et al., 2015) Marker selection was performed with CVRMS with 10-fold cross-validation, ten minimum markers, 20,000 maximum markers, 0.001 delta, and 0.8 accuracy cutoff, resulting in 1,530 marker subsets. The average correlation coefficient was 0.8001 (0.0705 for whole markers and 0.7743 for validation set). A scatterplot of predicted GEBV and observed values and of estimated fixed effects (β) and random effects (u) was generated (Supplementary Figure 2 and Supplementary Table 2). Since our method also considers the correlation between markers, we ensured that the selected markers were distributed genome-wide (Supplementary Figure 3). We compared the selected markers from CVRMS with those by the conventional top-down method. In the case of flowering time at the two locations, approximately twice as many markers should be selected in the top-down method to achieve the accuracy desired for the CVRMS results (Supplementary Figure 3).

CVRMS is a new method with a profound impact on the extracts of a trait-associated marker subset. Our method shows a high level of prediction accuracy for the real dataset using fewer markers than previous research regardless of the level of heritability. This tool is designed to select high-effect markers first, but correlative markers are excluded to minimize the loss of effectiveness and increase the accuracy of genomic prediction. Additionally, many iterations resulted in the elimination of splitting data-specific markers for increasing reproducibility.

## Methods

### Ridge regression

The use of ridge regression (RR) for GP was first proposed by Whittaker (Whittaker et al., 2000). RR minimizes the loss function, including the sum of squared regression. It is similar to the general least-square method, but different terms are added in which the loss function includes a model complexity measured by a sum-of-squares regression weight times a positive penalty parameter λ. This penalty prevents the overfitting from setting the weights to zero and makes the estimator a solution even in the case of multicollinearity and P≫N problems. Formally, we set N and P, the number of individuals and SNPs, respectfully. Let **y** be an outcome N × 1 vector, and **X** be a genotype N × P matrix of SNPs. A linear model is given by

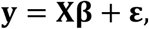

where **β** is the SNP effects vector, and **ε** is the error term of the linear model. The RR estimator minimizes

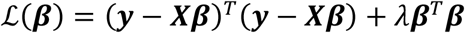

Using this loss function, the RR estimator of **β** is given by

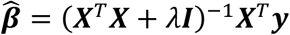

If λ is zero, RR corresponds to ordinary least squares (OLS). The OLS estimator exists only when rank(**X**^T^***X***) = ***P***. The OLS does not apply to the general GWAS situation (P≫N), but the RR estimator always has a solution if *λ* > 0 (Vlamming et al. 2015).

### Random forest

Random forests (RF) were first introduced by Breiman, which extended previous research into a random decision tree (Ho, 1995) and bagging of predictors (Breiman, 1996). RF constructs a forest of classification or regression trees that recursively divides groups of explanatory variables to predict the value of continuous response variables, such as human height, or of a categorical response, such as disease states. The algorithm of RF first randomly replaces a subset of samples to form a training dataset, and the resulting tree grows. Then, RF randomly selects a subset of predictive variables and retrieves the predictor that best partitions the response variables from the training dataset. Partitioning is based on identifying predictive variables that minimize the regression error (continuous response variable) or the within-group variance (categorical response variable). The optimal predictor is the first node in the tree and separates the data. For each node, a subset of predictive variables is randomly selected and continues this process until a full tree is grown. The power of the regression tree based on continuous variables is determined by the rate of variation of the response variables described by the tree. The importance of each predictive variable is estimated by the predictive power change of the tree after the predictive variable is randomly reordered among individuals. The predictive power of the tree is reduced when important features are permuted, but variables that affect the response variables less do not significantly affect the accuracy of the tree. As the trees become a forest, the importance value of the predictive variables is averaged across trees and analyzed to determine the value that best explains the variation of the response variable.

### Cross-validated Rank-based Marker selection

CVRMS requires three input files, the genotype, the phenotype, and a ranked marker table with GWAS p-values or marker effects. The form of the genotype file is a matrix, where the rows contain the samples, the columns have the markers, and a component is coded as a nonzero integer. The phenotype file is similar to the genotype file, but the columns contain traits instead of markers. The ranked marker table consists of the name of the SNP, chromosome and position, a p-value from the GWAS or marker effect size. The marker selection scheme of CVRMS is as follows (Figure 1):

1. Extract the validation set from input samples with a ratio of 0.1 and divide the remaining samples into k groups for cross-validation.
2. For each unique group:

A. Take the group as a holdout or test dataset
B. Take the remaining groups as a training dataset
C. Calculate the genomic estimated breeding value (GEBV) from the training set and the prediction accuracy using the test set for all markers
D. Retain the evaluation score and discard the model
E. Select the top two markers from the ranked marker set
F. Fit the model for the selected markers in the training set and the evaluation in the test set
G. If the rejection conditions (accuracy, maximum number of markers, delta, and time) are not satisfied, add the next marker and return to step (2-F)
3. CVRMS selects the final marker subset that is most frequently chosen for each group in step (2)
4. A final statistical model for genomic prediction is fitted for the final marker subset

**Figure 1.**
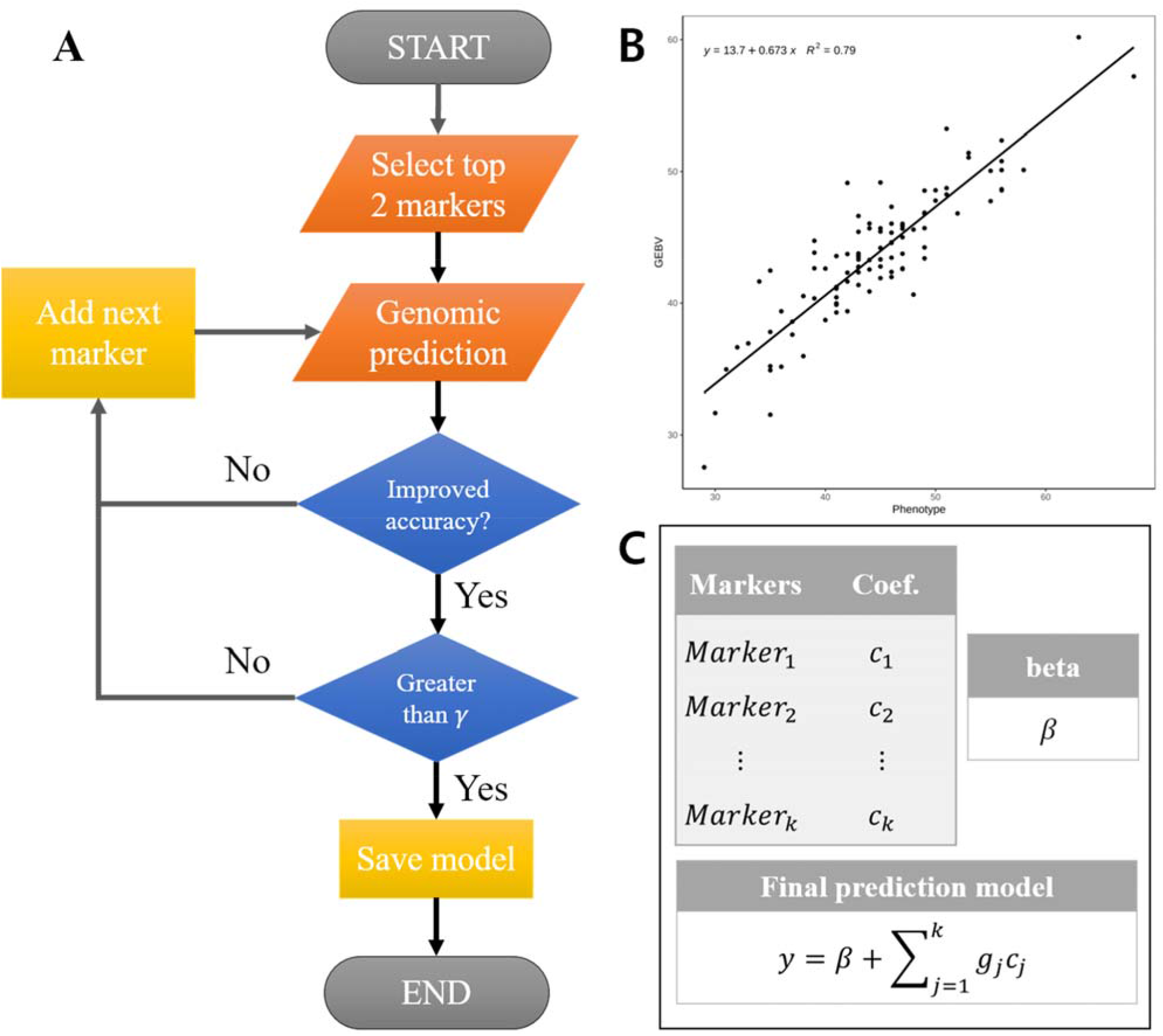
(A) Workflow, (B) scatterplot example, and (C) returned model. (A) is the marker selection workflow chart. (B) CVRMS provides a scatterplot of predicted GEBV and observed values and the linear regression line. (C) Coefficients for selected markers and β results are given as average values of all replicates.

This process considers the SNP-SNP interaction and increases the prediction accuracy by removing overlapping effects for SNPs, sampling bias, and low heritability problems. Therefore, our method can increase heritability and improve prediction accuracy by using fewer markers than choosing the upper SNPs from GWAS results. (Supplementary Figure ?) Additionally, CVRMS increases flexibility by allowing users to select the cut-off for the accuracy, the minimum and the maximum number of selected markers, increasing rate, time limits, and the fold of cross-validation.

## Supplementary Information

**Supplementary Table 1 Selected maker information table. Marker name in the input genotype array, chromosome, and position (base-pair) are included. CVRMS chooses markers for genomic prediction throughout the genome, rather than only from specific loci or gene that are significant in the association analysis.**

**Supplementary Table 2 CVRMS provide the result of marker selection with the marker information, prediction model including coefficient and β. To predict new data, multiply the new dataset by the coefficients and add beta in the prediction model to obtain the predicted value.**

**Supplementary figure 1.**
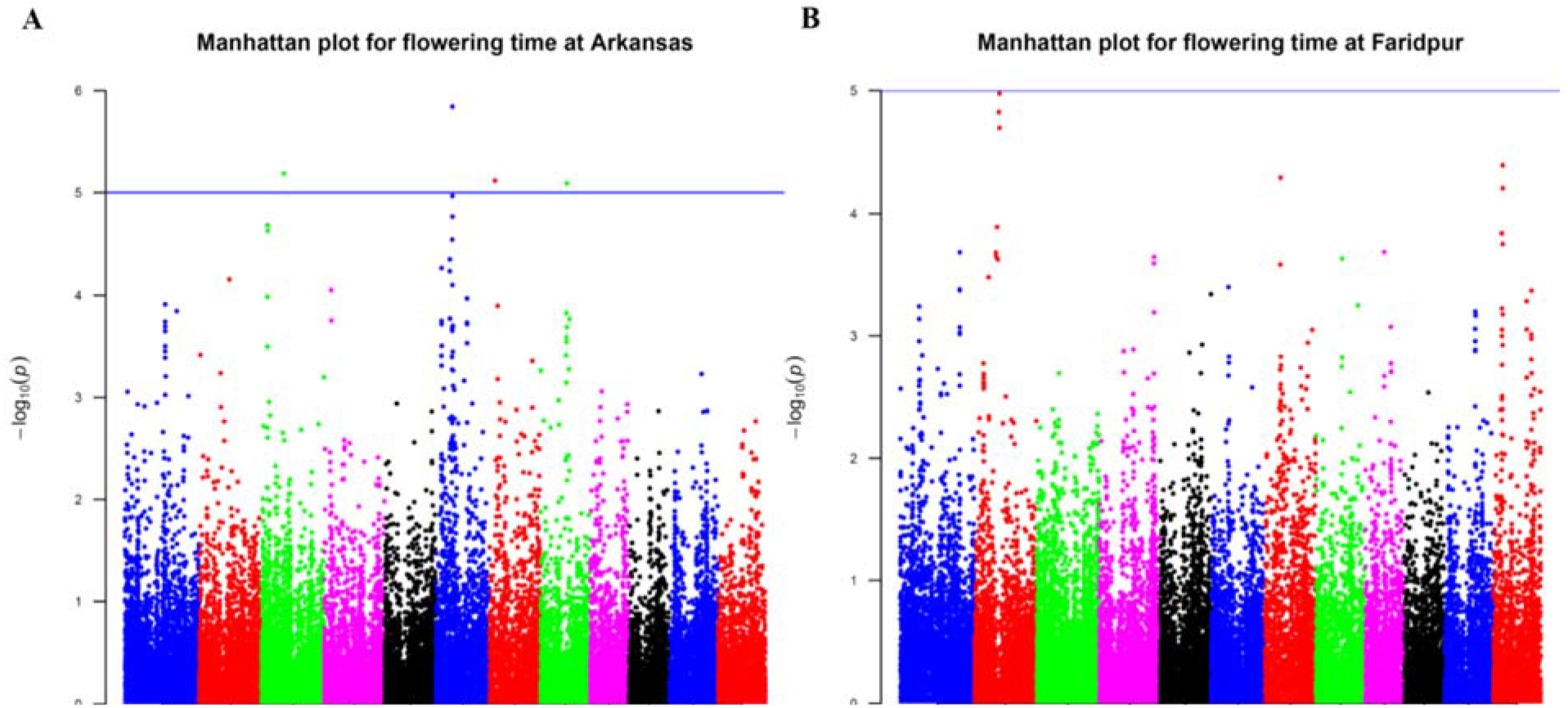
Manhattan plot for days to flowering at Arkansas (A) and Faridpur (B) from the result of GWAS (GAPIT version 2). P-value distributions are quite different and there is no significant SNPs in the result of Faridpur (B).

**Supplementary figure 2.**
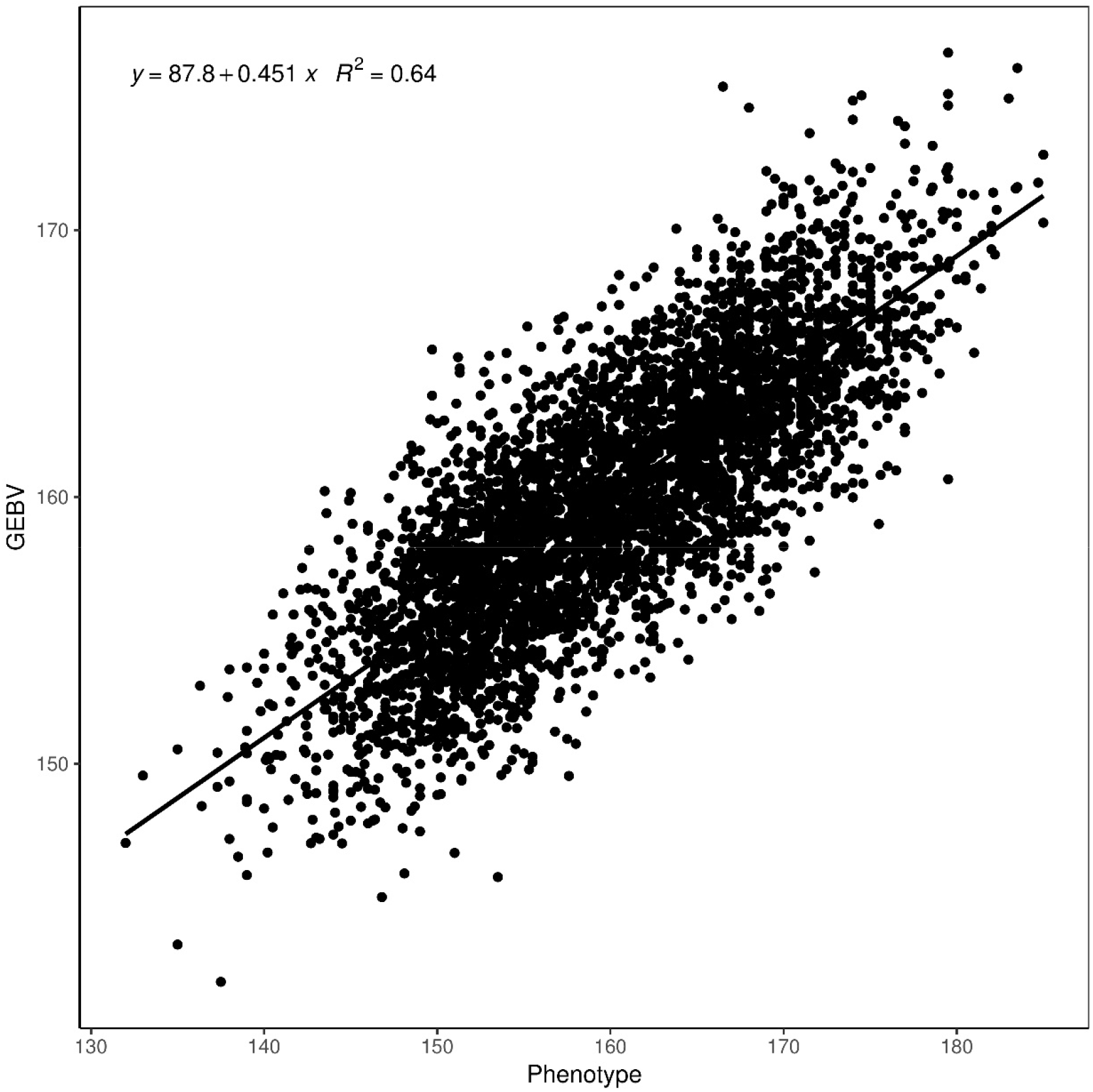
Scatter plot of the predicted and observed phenotypes. The estimated linear regression equation is provided top-left. Strong positive correlation is obtained and CVRMS provides relatively accurate prediction values.

**Supplementary figure 3.**
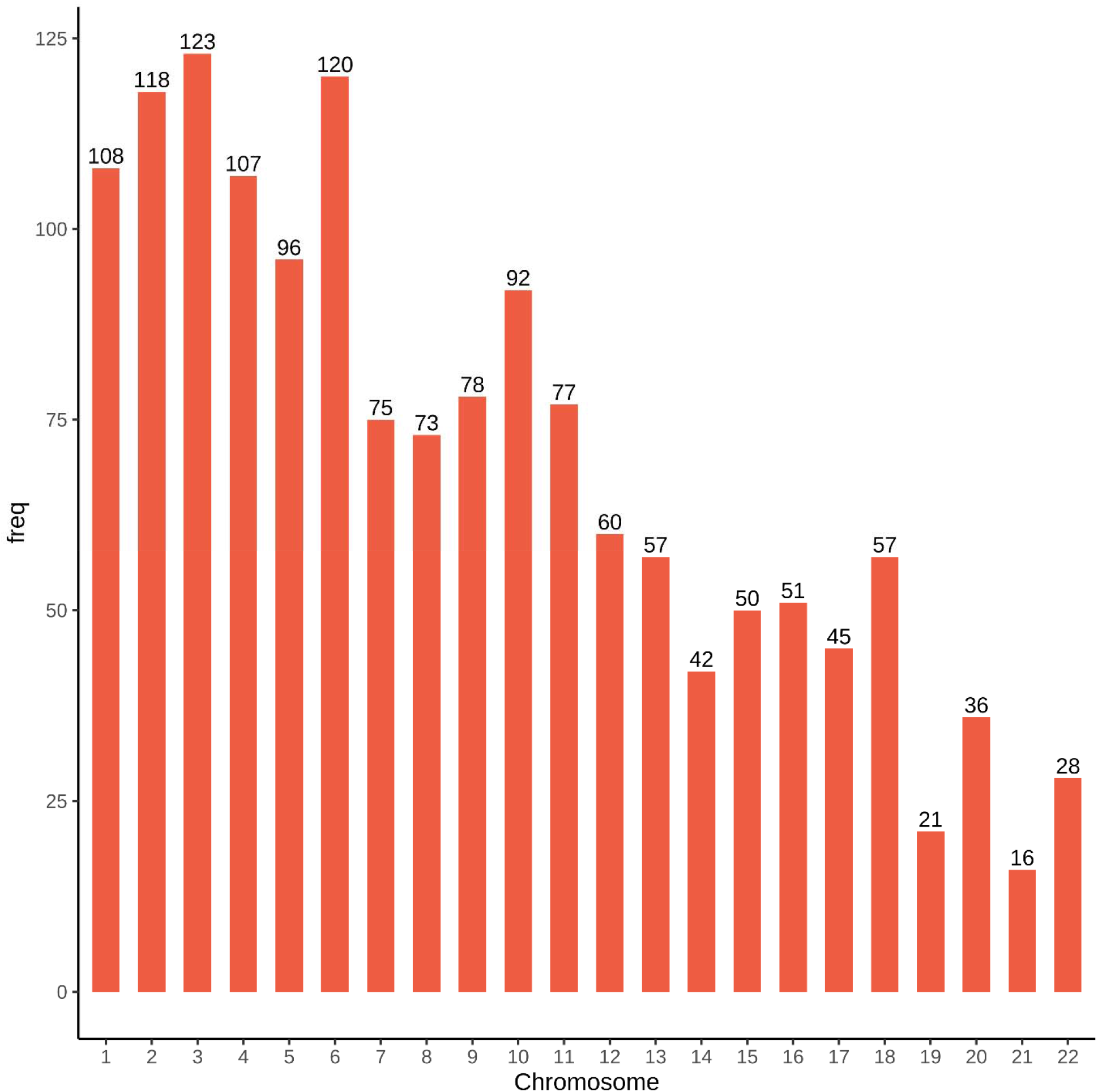
Chromosome distribution of selected markers. Genome-wide selection reduces bias rather than selecting only specific regions, e.g., significant SNPs in GWAS result.

